# Conserved structural features of RNA export pores spanning the double membrane of arterivirus and coronavirus replication organelles

**DOI:** 10.64898/2026.06.05.730354

**Authors:** Nina L. de Beijer, R.W.A.L. Limpens, Montserrat Bárcena, E.J. Snijder

**Affiliations:** Molecular Virology Laboratory, Leiden University Center of Infectious Diseases, Leiden University Medical Center, Leiden, Netherlands; Section Electron Microscopy, Department of Cell and Chemical Biology, Leiden University Medical Center, Leiden, Netherlands

**Author notes:** Corresponding author., Molecular Virology Laboratory, Leiden University Center of Infectious Diseases, Leiden University Medical Center, 2333 ZA Leiden, Netherlands.

## Abstract

Corona- and arteriviruses are distantly related positive-stranded RNA virus families within the order *Nidovirales*. Both transform intracellular membranes into double-membrane vesicles (DMVs) that serve as replication organelles. Newly made viral RNA presumably exits DMVs through double-membrane-spanning molecular pores formed by coronavirus nsp3-nsp4 and arterivirus nsp2-nsp3. Although sub-nm resolution structural information is available for the coronavirus DMV pore, only low-resolution data exist for the pore of the prototypic arterivirus equine arteritis virus (EAV). Here, we modeled arterivirus nsp2 and nsp3 to define their domain architecture, membrane topology, and potential organization into a DMV pore containing 12 copies of each protein. Despite the much smaller dimensions of the arterivirus pore, our analysis suggests striking conservation of features also found in coronaviruses, including conserved positively-charged residues lining analogous pore channel constrictions implicated in RNA export. We investigated their importance in arteriviruses using EAV reverse genetics and nsp2-nsp3 expression systems. While none of these mutations impaired DMV formation, charge-neutralizing substitutions were lethal, whereas conservative substitutions were partially tolerated. We also identified an additional conserved positively-charged residue in the nsp2 C-terminal domain, likely facing the outer DMV membrane, that is essential for DMV formation. Together, these findings reveal nidovirus-wide conservation of important DMV pore features despite the large evolutionary distance between arteri- and coronaviruses.

**Importance:** For eukaryotic positive-strand RNA viruses, efficient replication depends on the spatial organization of viral RNA synthesis within replication organelles in the infected cell’s cytoplasm. These virus-induced, membrane-bound compartments are thought to provide an optimized environment for viral RNA synthesis. In corona- and arteriviruses, distantly related families within the nidovirus order, unusual double-membrane vesicles (DMVs) have been identified as the primary site of viral RNA synthesis. A hallmark of these structures is a double-membrane-spanning pore complex that connects their interior to the cytosol, thereby presumably enabling export of newly synthesized viral RNA. Here, we analyze the arterivirus DMV pore complex and identify several conserved structural features, including positively-charged residues in the pore-forming proteins nsp2 and nsp3, which are essential for viral replication, likely due to their role in RNA export. Together, our findings underscore the functional conservation and fundamental importance of DMV pore complexes across evolutionary distant nidovirus families.

## Introduction

*Nidovirales*, a rapidly expanding order of positive-stranded RNA viruses, currently comprises 14 families and includes numerous human and veterinary pathogens (1, 2). Members of this order display remarkable diversity in both genome size and host range. Recent metagenomic studies have further broadened this diversity, revealing increasing variation in genome size and organization, including invertebrate nidovirus genomes exceeding 60 kilobases (3) and the first examples of segmented nidovirus genomes (4). Despite their diversity, nidoviruses share a broadly similar genome organization and encode a conserved array of core replicative enzymes that include the viral RNA polymerase (3–6). Their activity is supported by additional viral nonstructural proteins (nsps), which perform diverse functions and display varying degrees of conservation across the order.

Among nidovirus families, the *Coronaviridae* has been studied most extensively as it includes important human pathogens such as severe acute respiratory syndrome coronavirus (SARS-CoV) and SARS-CoV-2. The family *Arteriviridae* has likewise been characterized in considerable detail given the veterinary importance of several members, including porcine reproductive and respiratory syndrome virus (PRRSV) and equine arteritis virus (EAV). Notably, among nidoviruses infecting vertebrate hosts, arteriviruses represent the most distantly related lineage to coronaviruses. Their genomes are approximately half the size of those of coronaviruses, highlighting the substantial evolutionary divergence between these two nidovirus families (4, 7). In both corona- and arteriviruses, a large replicase gene comprising open reading frames (ORFs) 1a and 1b occupies more than two-thirds of the 5’-proximal part of the genome. It encodes the nonstructural polyproteins pp1a and pp1ab, the latter being produced via a programmed ribosomal frameshift occurring just upstream of the ORF1a stop codon. Two to five ORF1a-encoded protease domains mediate the co- and post-translational cleavage of pp1a and pp1ab to release 13 to 16 mature nsps that drive viral genome replication and subgenomic mRNA synthesis, in addition to engaging in a range of virus-host interactions (8, 9). Besides encoding components of the viral replication and transcription complex, pp1a and pp1ab also include three nsps containing multi-spanning transmembrane domains (TMDs). In coronaviruses, these are nsp3, nsp4, and nsp6, which correspond to nsp2, nsp3, and nsp5 in the arteriviruses (Figure 1A). In the infected cell, these proteins transform endoplasmic reticulum (ER)-derived membranes into specialized replication organelles (ROs), including numerous double-membrane vesicles (DMVs), whose interior is believed to serve as the primary site of viral RNA synthesis (10–14).

**Figure 1:**
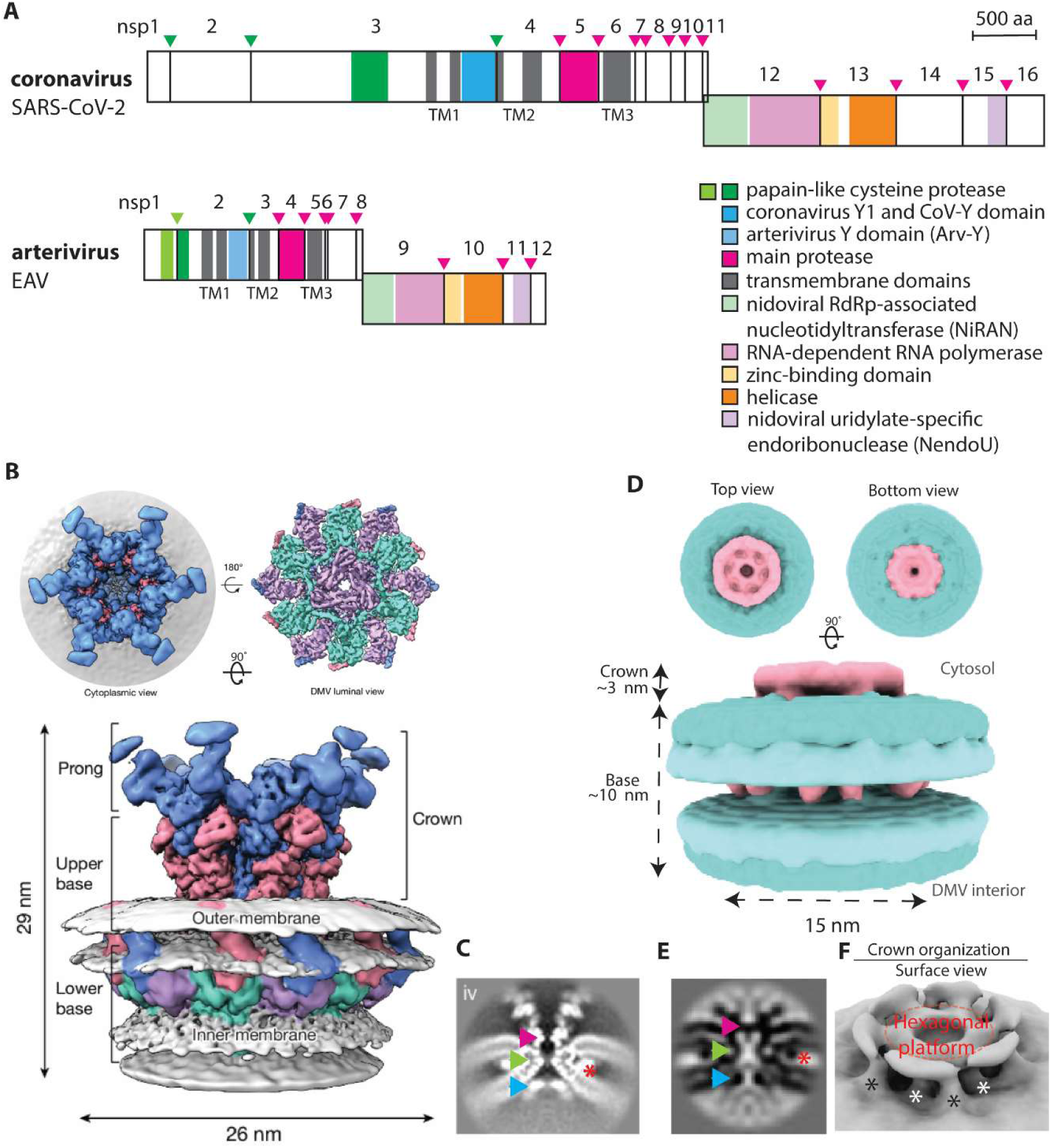
Comparison of arteri- and coronavirus replicase polyproteins and DMV pore complex structures. (A). Schematic overview of replicase pp1ab of coronaviruses (SARS-CoV-2) and arteriviruses (EAV). A ribosomal frameshift signal just upstream of the ORF1a stop codon directs the extension of the ORF1a-encoded polyprotein (pp1a) to produce pp1ab. ORF1a-encoded transmembrane domains (TMDs) are indicated in grey, whereas proteases and their corresponding polyprotein cleavage sites are indicated by arrows in matching colors. ORF1a of both families encodes three transmembrane domain-containing proteins that flank the protease domains, whereas ORF1b encodes several conserved domains involved in viral RNA synthesis (see panel A legend). (B) Subtomogram averaging map for pore complexes in coronavirus DMVs isolated from a SARS-CoV-2 nsp3-nsp4 expression system (4.9 Å overall resolution), with nsp3 depicted in pink and blue and nsp4 in cyan and purple (17). (C) Cross-section through the density map of the SARS-CoV-2 nsp3-nsp4 pore channel. Three constrictions (colored arrowheads: magenta, green, blue) and the position of the interacting luminal domains of nsp3 and nsp4 (red asterisk) are indicated. (D) Low-resolution (20 Å) subtomogram averaging map of the arterivirus pore complex obtained by in situ cryo-electron tomography of EAV-infected cells (25). (E) Three putative constriction sites along the EAV DMV pore channel are indicated by colored arrowheads; the putative position of the interacting luminal domains of EAV nsp2 and nsp3 is labeled with a red asterisk. (F) Surface view of the crown of the EAV pore complex showing a hexagonal platform surrounding the cytosolic opening of the pore channel. Two sets of putative connectors emerge from the pore’s upper base and connect to the small crown of the pore: the inner stalks (white asterisks) and outer stalks (black asterisks).

The ultrastructure of coronavirus- and arterivirus-induced DMVs has been examined in considerable detail in recent studies using in situ cryo-electron tomography of infected cells. In coronaviruses, these studies revealed an nsp3-containing molecular pore complex with sixfold symmetry that spans the DMV’s double membrane (15). As DMVs had long been assumed to be fully sealed, the identification of this putative RNA export channel resolved a key question regarding how newly synthesized viral RNA may be transported from the DMV interior to the cytosol, where translation and encapsidation occur. Subsequent studies revealed that co-expression of SARS-CoV-2 nsp3 and nsp4 suffices to induce the formation of both DMV and pore complexes that mimic those observed in infected cells (16). More recently, using DMVs purified from an nsp3-nsp4 expression system lacking viral RNA synthesis, the structure of a nsp3-nsp4 pore complex was resolved at unprecedented resolution by cryo-EM and subtomogram averaging (17). This study revealed a distinct stoichiometry and architecture, which consists of two stacked rings of six nsp3 molecules and two rings of six nsp4 molecules, together forming a sixfold-symmetric complex surrounding a trans-membrane channel (Fig. 1B). The nsp3 rings reside in the outer DMV membrane, forming the upper base and crown of the pore complex, whereas the nsp4 rings constitute the lower base facing the DMV interior. Within the DMV intermembrane space, the 12 copies of nsp3 and nsp4 interact through their luminal domains, forming a belt-like structure encircling the complex. Along the coronavirus pore channel, rings of positively-charged amino acid residues, which are highly conserved among betacoronaviruses, mark three constriction sites (Fig. 1C; (17)). Their positioning within the channel provides additional support for the pore’s proposed role in RNA export, as protein-RNA interactions are frequently mediated by electrostatic interactions (18, 19). Charge-neutralizing or charge-reversing mutations of these channel-facing residues yielded nonviable SARS-CoV-2 mutants (17). Notably, for two of the three constriction sites these mutants could be rescued by introducing an alternative positively-charged residue. Studies using an nsp3-nsp4 expression system indicated that these mutations did not affect DMV morphology or nsp3-nsp4 interactions, supporting a role for the conserved charges in pore function rather than DMV biogenesis (17). Although arterivirus DMVs are considerably smaller than those induced by coronaviruses (20), a striking parallel is that their biogenesis can be induced by co-expression of nsp2 and nsp3, the arterivirus counterparts of coronavirus nsp3 and nsp4 (20–24).

Recently, we employed in situ cryo-electron tomography to visualize arterivirus DMV pores for the first time in cells infected with PRSSV and EAV, as well as in cells co-expressing EAV nsp2 and nsp3 (25). Subtomogram averaging revealed a sixfold-symmetric EAV pore complex measuring ∼13 nm in height and ∼15 nm in width (Fig. 1D). Its central channel spans both DMV membranes and, as in coronavirus DMV pores, contains three constrictions. Notably, the middle constriction appears to be encircled by 12 EM densities that were postulated to represent pairs of interacting ER-luminal domains of EAV nsp2 and 3 (Fig. 1E) (25), resembling the architecture of the SARS-CoV-2 nsp3-nsp4 pore (17). Despite these similarities, the arterivirus pore-forming proteins are substantially smaller than their coronavirus nsp3 and nsp4 counterparts (Figure 1A), reflecting the considerable evolutionary distance between these nidovirus families and likely explaining the absence of the prominent cytosolic crown that is characteristic of coronavirus DMV pores. Nevertheless, a smaller sixfold symmetric crown is present on the cytosolic side of the arterivirus DMV pore. Particularly prominent is the hexagonal platform situated directly above the upper constriction site, which was postulated to be formed by the conserved C-terminal domain of nsp2 (25). In analogy with coronavirus nsp3 domain nomenclature (17), we will refer to this region as the arterivirus Y domain (Arv-Y; Fig. 1a; see also below). Furthermore, two distinct sets of stalks connect the upper base of the arterivirus pore to the cytosolic crown. This may reflect a folding arrangement similar to that observed in the coronavirus pore complex, which would involve an interaction between the N-terminal (PLP2) and Arv-Y domains of arterivirus nsp2 (Fig. 1F) (17, 25). We have now analyzed available arterivirus nsp2-nsp3 sequences in detail and modeled components of a putative DMV pore complex, revealing several similarities to the coronavirus DMV pore architecture. Notably, the arterivirus DMV pore also appears to harbor rings of positively-charged residues positioned at three constrictions along the pore channel. Alanine substitution of these residues abolished EAV replication, whereas replacement with alternative positively-charged residues was tolerated to some extent. Collectively, our data indicate that key structural and functional features of the DMV pore complex have been conserved across the vast evolutionary distance separating the corona- and arterivirus branches of the nidovirus order.

## Methods

### Sequence alignments and structural predictions

To perform multiple-sequence alignments (MSA) of arterivirus nsp2 and nsp3, the 23 available reference genomes listed under *Arteriviridae* in the master species list (MSL) 39v2 [ICTV] were downloaded from GenBank and included for further analysis. After translating ORF1a, the nsp2 sequence was extracted starting from the N-terminal border of the PLP2 domain, since any upstream residues were not expected to be part of the core of the pore complex, up to the (predicted) nsp2/3 cleavage site (Fig. S1) (26). For nsp3, the full-length sequence between the (predicted) nsp2/3 and nsp3/4 cleavage sites was selected (Fig. S2). The GenBank accession numbers used for the alignments and abbreviations used in subsequent figures are listed in Supplementary Table 1. Nsp2 and nsp3 sequences were aligned separately using Clustal Omega (27) and analyzed in Jalview2.11.5.0. WebLogo3.0 (28) was used to create sequence logos; complete alignments can be found in Fig. S3 (nsp2) and S4 (nsp3).

AlphaFold3 (29) was used to predict protein structures using sequences translated from the reference genomes described above. The input sequences used for each of the predictions shown in this paper can be found in Supplementary Table 2. PyMOL (Molecular Graphics System, Version 3.0.3 Schrödinger, LLC) was used to annotate and visualize the residues of interest and find residues potentially involved in chain-chain interactions (find>polar interactions). The predicted Local Distance Difference Test (pLDDT) plots were exported from PyMOL and visualized in GraphPad Prism (version 10.6.1). The EAV-Y1/Y2 hexamer was placed in the density map (25) with a weighted threshold of 1 using the fit-in-map tool in ChimeraX (30).

### EAV reverse genetics

EAV mutants were generated by mutagenesis of full-length cDNA clone pEAN551 (31, 32) with QuickChange using ACCUZYME DNA Polymerase [Meridan] and the primers listed in Supplementary Table 3. Whole-plasmid sequencing was performed at Plasmidsaurus using Oxford Nanopore Technology, with custom analysis and annotation. Full-length cDNA clones were linearized using *Xho*I [NEB] and transcribed into RNA using the *in vitro* T7 RNA polymerase transcription kit [Thermo Fisher Scientific]. For each mutant, 5 μg of RNA was transfected into 5 × 10^6^ baby hamster kidney cells (BHK-21; ATCC CCL10) using the Amaxa Cell Line Nucleofector kit T [Lonza] and program A-031. Cells were incubated at 37°C, supernatant samples were harvested at 10 , 24 , 48, and 120 h after the launch, and progeny virus titers were determined via plaque assay on BHK-21 cells. Plaque assays were essentially performed as previously described (33), but at 37°C instead of 39.5°C. Viral RNA was isolated from the harvested supernatants using TriPure [Roche] according to the manufacturer’s instructions and sequenced using Sanger sequencing after RT-PCR amplification of specific fragments with the primers listed in Supplementary Table 4.

### EAV nsp2-3 expression and immunoprecipitation

Plasmids used for the CMV promoter-driven expression of EAV nsp2 or nsp3 mutants were based on pcDNA3.1-EAV-HAnsp23GFP (20). This construct encodes a self-cleaving nsp2-3 polyprotein, yielding N-terminally HA-tagged nsp2 and C-terminally GFP-tagged nsp3. NCl-H1299 cells (kindly provided by Dr. Marc Vooijs, Maastricht University) were transfected with wild-type or mutant EAV nsp2-3 expression constructs using Lipofectamine 2000 [Invitrogen] according to the manufacturer’s protocol. The pcDNA3.1 empty vector was included as a negative control. After 24 h, cell lysates were harvested using lysis buffer (10 mM Tris-HCl (pH 7.5), 150 mM NaCl, 0.5 mM EDTA, 1% Triton X-100). All buffers used during co-immunoprecipitation (co-IP) experiments were supplemented with cOmplete protease inhibitor [Roche]. Nuclei were removed by low-speed centrifugation and the protein concentration of the post-nuclear supernatant (PNS) was determined using a BCA assay [Pierce]. Equal amounts of protein were used as input for (co)immunoprecipitation (IP) analysis. GFP-catcher beads [Antibodies online] were equilibrated in washing buffer (10 mM Tris-HCl (pH 7.5), 150 mM NaCl, 0.5 mM EDTA). After PNS was added to the equilibrated beads, the mixture was incubated for at least 1 h at 4°C with constant agitation. The beads were centrifuged at 2,700 g for 2 min at 4°C and washed, first with washing buffer and subsequently with a high-salt washing buffer (10 mM Tris-HCl (pH 7.5), 300 mM NaCl, 0.5 mM EDTA). To elute proteins bound, beads were boiled in 2x Laemmli sample buffer for 10 min. The samples were run on a 10% sodium dodecyl sulfate-polyacrylamide (SDS-PAGE) gel and transferred to a low-fluorescence PVDF membrane using a Trans-blot Turbo system [Bio-Rad]. Membranes were blocked for 1 h at room temperature using 1% casein in PBS with 0.05% Tween-20. The following primary antibodies were used for western blot detection: rabbit anti-GFP [042150, LUMC (34)], rabbit anti-GAPDH [clone 14C10, Cell Signaling Technology], and rabbit anti-EAV-nsp2-CTD. The latter was kindly provided by Dr. Marjolein Kikkert (LUMC) and was generated by a commercial service provider (GenScript) using a standard immunization protocol and a combination of KLH-coupled synthetic peptides derived from the C-terminal domain of EAV nsp2: CASTVDPHSFDQKK and CGDFLKLNPGFRLIGG. The reactivity and specificity of the antiserum was previously confirmed using EAV-infected cells, Western blotting and immunofluorescence microscopy. As secondary antibodies for western blot detection, donkey anti-rabbit-Cy3 [711-165-152, Jackson] was used for the co-IP samples and swine anti-rabbit-HRP [P-0217, DAKO] for the input samples. The blots of the input samples were visualized using Clarity Western ECL substrate [Bio-Rad]. All blots were visualized using an Alliance Q9 advanced imaging system [Uvitec].

### Transmission electron microscopy

For transmission electron microscopy (TEM), NCl-H1299 cells were transfected with the nsp2-3 expression constructs as described above. After 24 h, the cells were fixed using 1.5% glutaraldehyde in 0.1 M cacodylate buffer (pH 7.4) for 1 h and stored at 4°C in 0.5% paraformaldehyde in 0.1 M PHEM until further processing. The cells were then stained for 1 hour at 4°C with 1% (wt/vol) OsO_4_ in 0.1 M cacodylate buffer. Hereafter, the cells were washed in 0.1 M cacodylate buffer and Milli-Q water (MQ) and stained for 1 h with 1% (wt/vol) uranyl acetate in MQ. Subsequently, the samples were dehydrated by incubation in increasing ethanol concentrations (from 70% to 100%) and embedded in epoxy resin [LX-112, Ladd Research], which was polymerized at 60°C. Next, the samples were sectioned using an ultra-microtome and collected on carbon-coated mesh-100 copper EM grids. The sections were post-stained with 7% (wt/vol) uranyl acetate and Reynold’s lead citrate. Images were collected in a Tecnai12 Twin transmission electron microscope equipped with a OneView 4k high-frame rate CMOS camera (Gatan).

### Statistical analysis and software

The experimental data in this paper are represented as mean ± standard deviation, and statistical significance was calculated using the unpaired Student’s t-test. All graphs and statistical calculations were made in Prism version 10 [GraphPad], and the significance is represented by asterisks as follows: P<0.05 = *, P<0.01 = **, P<0.001 = ***, P<0.0001 = ****. The figures and schematics were created using Adobe Illustrator 2025.

## Results

### Domain organization and membrane topology of arterivirus nsp2 and nsp3

Given the broad similarities between coronaviruses and arteriviruses in DMV formation and pore complex architecture (25), we investigated the extent to which key viral proteins involved in these processes share sequence similarities that could translate into common structural features. Specifically, we focused on arterivirus nsp2 and nsp3, which constitute the minimal machinery required to form DMV-spanning pore complexes (25), analogous to their coronavirus counterparts nsp3 and nsp4 (16). We first used Clustal Omega to align the nsp2 and nsp3 sequences from all 23 currently recognized arterivirus species, as defined in the ICTV master species list (MSL; version 41 (1)) (Fig. S3-S4). For nsp2, sequences were aligned from the N-terminal boundary of the papain-like protease 2 (PLP2) domain (35) to the (predicted) nsp2/3 cleavage site, whereas full-length nsp3 sequences were used. The resulting alignments were evaluated for consistency with previously reported features, including predicted transmembrane regions (36) and the experimentally validated ER-luminal domain of nsp3, which contains a cluster of conserved cysteines (32). Despite substantial differences in overall length, our analysis revealed a broadly similar organization among arterivirus nsp2 proteins (Fig. 2). Specifically, the highly conserved N-terminal PLP2 domain is followed by a hypervariable region (HVR) ranging from approximately 100 to 700 residues, a predicted ER-luminal domain flanked by two transmembrane regions (TMH1-3 and TMH4, see also below), and a moderately conserved C-terminal domain that is predicted to be cytosolic (Fig. 2A) to ensure PLP2 access to the nsp2/3 cleavage site. This C-terminal region of nsp2 was recently described to comprise not only the Y1 domain, which is conserved across the nidovirus order, but also a Y2 domain, which is conserved in most vertebrate-infecting nidovirus families, and several structural features of a Y4 domain, which is found in coronaviruses (37). For simplicity, we will refer to this Y1-Y2-Y4 region as the arterivirus Y (Arv-Y) domain throughout this paper.

**Figure 2:**
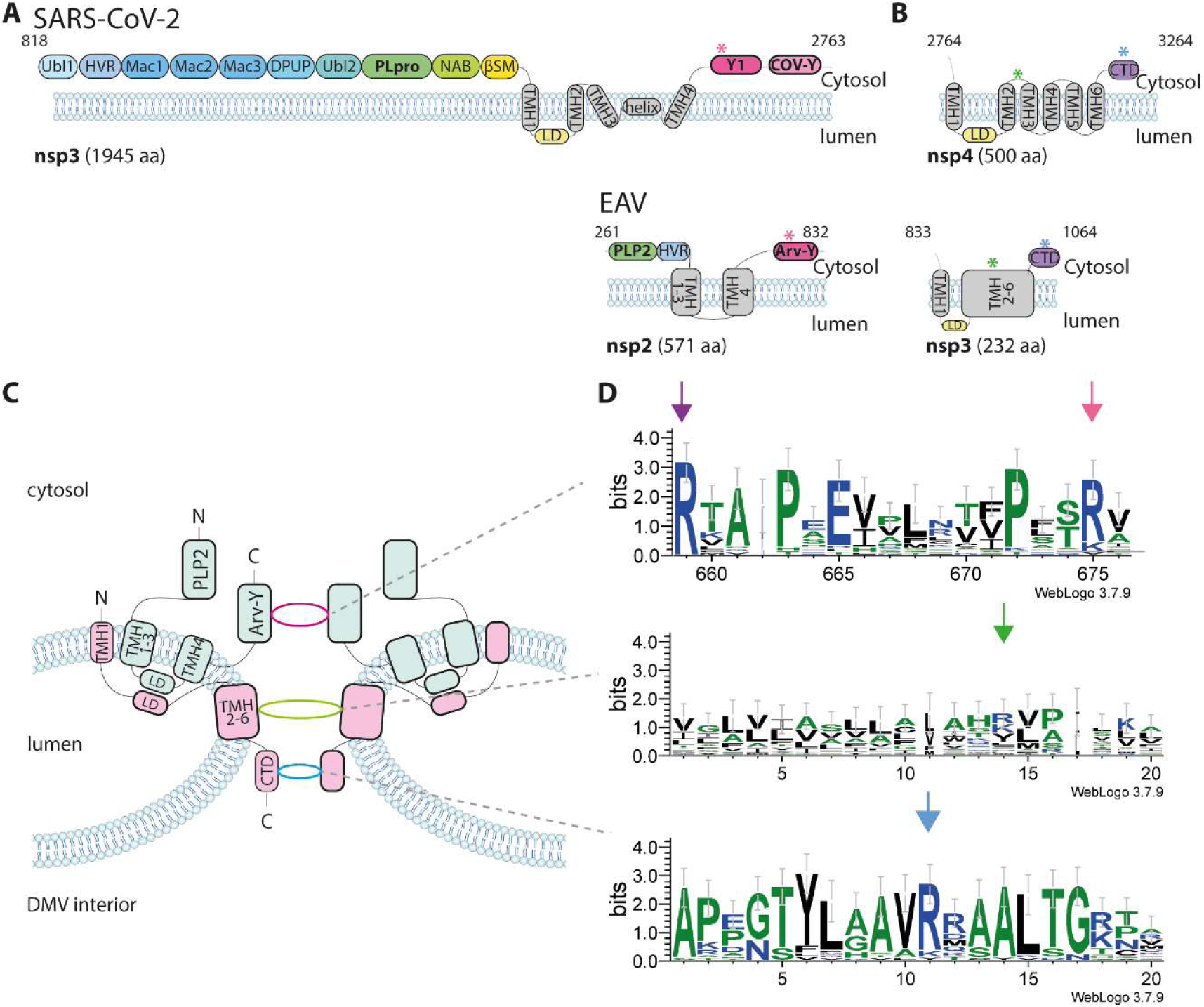
A model of the organization of the arterivirus DMV pore complex and the conservation of positively-charged residues lining the pore channel. (A) The overall domain organization of SARS-CoV-2 nsp3, as resolved previously (16, 17, 45), and EAV nsp2 is depicted, with the positions of the previously established positively-charged residues lining the three DMV pore constrictions marked with colored asterisks. (B) Domain organization of SARS-CoV-2 nsp4 (16, 17, 45) and EAV nsp3. The positions corresponding to the established are marked with colored asterisks. (C) Working model of the arterivirus DMV pore complex, consisting of two hexameric rings of nsp2 (cyan) and two hexameric rings of nsp3 (pink), with only two copies of each protein depicted for clarity. The model is based on arterivirus sequence comparisons and structural predictions, using the sub-nm-resolution structure available for the SARS-CoV-2 nsp3-nsp4 pore complex (17) as a template. (D) Identification of conserved positively-charged residues in arterivirus nsp2 and nsp3. Regions of interest from arterivirus-wide sequence comparisons are depicted using WebLogo 3.0 (28). The nsp2 Arv-Y domain predicted to line the first constriction (magenta oval) contains two conserved positively-charged residues (Fig S3; residues R659 and R673 in EAV pp1a). The nsp3 region proposed to face the second constriction (green oval) is located between TMH2 and TMH3 and contains a positively-charged residue that is conserved in about half of the arterivirus nsp3 sequences (R920 in EAV pp1a; green arrow). Most viruses lacking this conserved positive charge feature an aromatic amino acid (Fig. S4). The domain proposed to face the third constriction (blue oval) is located in the nsp3-CTD and contains a highly-conserved positive charge (Fig. S4; R1043 in EAV pp1a; blue arrow).

Our understanding of the biological function of the nsp2 HVR remains incomplete, but this domain has been implicated in viral fitness, virulence, and interactions with host factors and pathways. It likely serves as a flexible module enriched in intrinsically disordered segments (38, 39). The considerable size variation of the HVR accounts for most of the overall size differences observed among arterivirus nsp2 proteins, which range from 571 residues in EAV to 1201 residues in the rodent RtMruf-arterivirus. Its coronavirus counterpart, nsp3, is also characterized by substantial size variation and lineage-specific insertions and deletions (40); however, at approximately 2,000 residues in most family members, it is markedly larger than arterivirus nsp2. This contrast is particularly striking for EAV nsp2, which contains an HVR of just over 100 residues.

In contrast, arterivirus nsp3 exhibits comparatively limited size variation, ranging from 224 to 279 residues, and displays a more consistent domain architecture. Immediately downstream of the nsp2/3 cleavage site lies an N-terminal transmembrane helix (TMH), followed by a known ER-luminal domain that likely mediates the interaction with nsp2 (25, 32, 41). This arrangement would mirror that of the luminal domains of SARS-CoV-2 nsp3 and nsp4, whose interaction appears to be crucial for the formation of the DMV pore (17, 42). Downstream of the ER-luminal domain, nsp3 proteins from all arteriviruses contain a second transmembrane region predicted to comprise five TMHs (TMH2-6; see also below), followed by a cytosolic C-terminal domain (nsp3-CTD) (Fig. 2B). Overall, this topology, with both nsp3 termini located in the cytosol to ensure cleavage by PLP2 and the nsp4 main protease, closely resembles that described for coronavirus nsp4 (∼500 residues) (17, 43, 44), despite the arterivirus protein being about half the size of its coronavirus counterpart.

The shared molecular features of arterivirus nsp2-nsp3 and coronavirus nsp3-nsp4, together with their established roles in DMV and pore formation and likely similar stoichiometry within the DMV pore complex, led us to hypothesize that the arteri- and coronaviral pore complexes may share key architectural features. Specifically, as also proposed elsewhere (25), we assume that the sixfold-symmetric pore complex comprises two sets of six nsp2 molecules forming the upper base and cytosolic crown of the structure, with one ring directly facing the pore channel, and two sets of six nsp3 molecules forming the lower base of the complex. Based on these assumptions, we generated a working model of the domain organization and membrane topology of the arterivirus DMV pore complex, using the published sub-nanometer-resolution structure of the SARS-CoV-2 pore complex as a structural template (Fig. 2C) (17).

### AlphaFold modeling supports structural similarities between arterivirus nsp2-nsp3 and coronavirus nsp3-nsp4

As only a low-resolution model of the arterivirus DMV pore complex is currently available (25) (20 Å resolution), we used AlphaFold 3 to tentatively model the most likely components of the DMV pore complexes of EAV and PRRSV2, two phylogenetically distant members of the arterivirus family. The resulting predictions were evaluated using AlphaFold’s confidence scores and their consistency with established topological constraints, such as the ER-luminal localization of specific domains and the established cytosolic localization of the N- and C-termini of nsp2 and nsp3. During the finalization of this work, a detailed set of structural predictions for the TMD2-CTD region of coronavirus nsp3 and arterivirus nsp2 was published (37). These findings are taken into account in our discussion of data pertaining to this specific region.

We first modeled the structures of nsp2 and nsp3 monomers from both viruses (Fig. S5). For nsp2, well-resolved domains with a per-residue measure of local confidence (pLDDT) >70 included the cytosolic PLP2 and Arv-Y domains in both viruses, as well as the TMHs of PRRSV2 nsp2. Consistent with previous studies, the HVR located between PLP2 and the nsp2 TMD lacked a well-defined structural organization, resulting in low overall predicted template-modeling scores (pTM) (Fig. S5). The AlphaFold-derived model revealed that, within the predicted transmembrane regions, several α-helices were estimated to be only 12 residues or shorter, which is likely insufficient to span the lipid bilayer (46). This limitation complicates the accurate prediction of the exact conformation and membrane topology of the nsp2 TMD. However, given that both termini of the protein are expected to reside on the cytosolic side of the membrane, the total number of membrane crossings must be even. Accordingly, this transmembrane region most likely comprises four TMHs, along with an additional short helix preceding TMH4, which is too short to fully cross the membrane. The nsp2 Arv-Y domain includes the conserved Y1 and Y2 domains (see above), which contain several conserved cysteine residues. The preservation of both sequence and structural features of this region observed in our models across arteri- and coronaviruses is consistent with recently published work (37) and suggests a shared, yet uncharacterized, functional role for the Arv-Y domain.

For EAV nsp3, only the structure of the CTD was predicted with high confidence (pLDDT>70), whereas PRRSV2 nsp3 was modeled with higher overall confidence (pTM = 0.59), except for the second putative TMH and several short flexible regions. Overall, the structural organization was as expected, with an N-terminal TMH1, an ER-luminal domain, a cluster of five TMHs (TMH2-6), and the CTD. The arterivirus nsp3-CTD contains a conserved winged helix-turn-helix motif that is also present in the CTD of coronavirus nsp4 and has been proposed to mediate interactions with RNA (47). This motif was readily identifiable in the predicted structures of both EAV and PRRSV2 nsp3 (Fig. S5E,F). Overall, the AlphaFold monomer models were consistent with the domain organization presented in Fig. 2 and recapitulated the conserved structural features previously described for arterivirus nsp3 proteins.

Earlier experimental studies demonstrated a strong interaction between arterivirus nsp2 and nsp3, likely involving the nsp3 ER-luminal domain that is located between TMH1 and the remaining transmembrane regions (comprising TMH2-6) (32). However, the boundaries of the nsp2 luminal domain engaging in this interaction remained unknown. To address this, we modeled nsp2-nsp3 heterodimers for EAV and PRRSV (Fig. 3). For both viruses, predicted aligned error (PAE) plots indicated high confidence in the relative positioning of the nsp2 transmembrane regions and the nsp3 ER-luminal domain. In EAV, the nsp2 region predicted by AlphaFold to engage in the interaction with nsp3 corresponds to the segment between TMH1-3 and TMH4 in the sequence alignment (Fig. 2B; EAV pp1a residues N571-P602). For PRRSV2, this corresponds to pp1a residues P1311-P1350 (Fig. S3), which is consistent with the recent report that the transmembrane region of nsp2 is essential for the heterodimerization of PRRSV2 nsp2-nsp3 (48). Several plausible polar interactions (inter-residue distance < 3.5 Å) were identified between residues located within the predicted ER-luminal regions of nsp2 and nsp3 (Fig. 3). Across arteriviruses, the conserved residues in this predicted ER-luminal domain of nsp2 include two cysteines (Fig. 3C), while the established luminal domain of nsp3 contains four conserved cysteines (Fig. 3D). Both the predicted luminal domain of nsp2 and the tentative TMH annotation were incorporated into our model of the pore complex (Fig. 2C).

**Figure 3:**
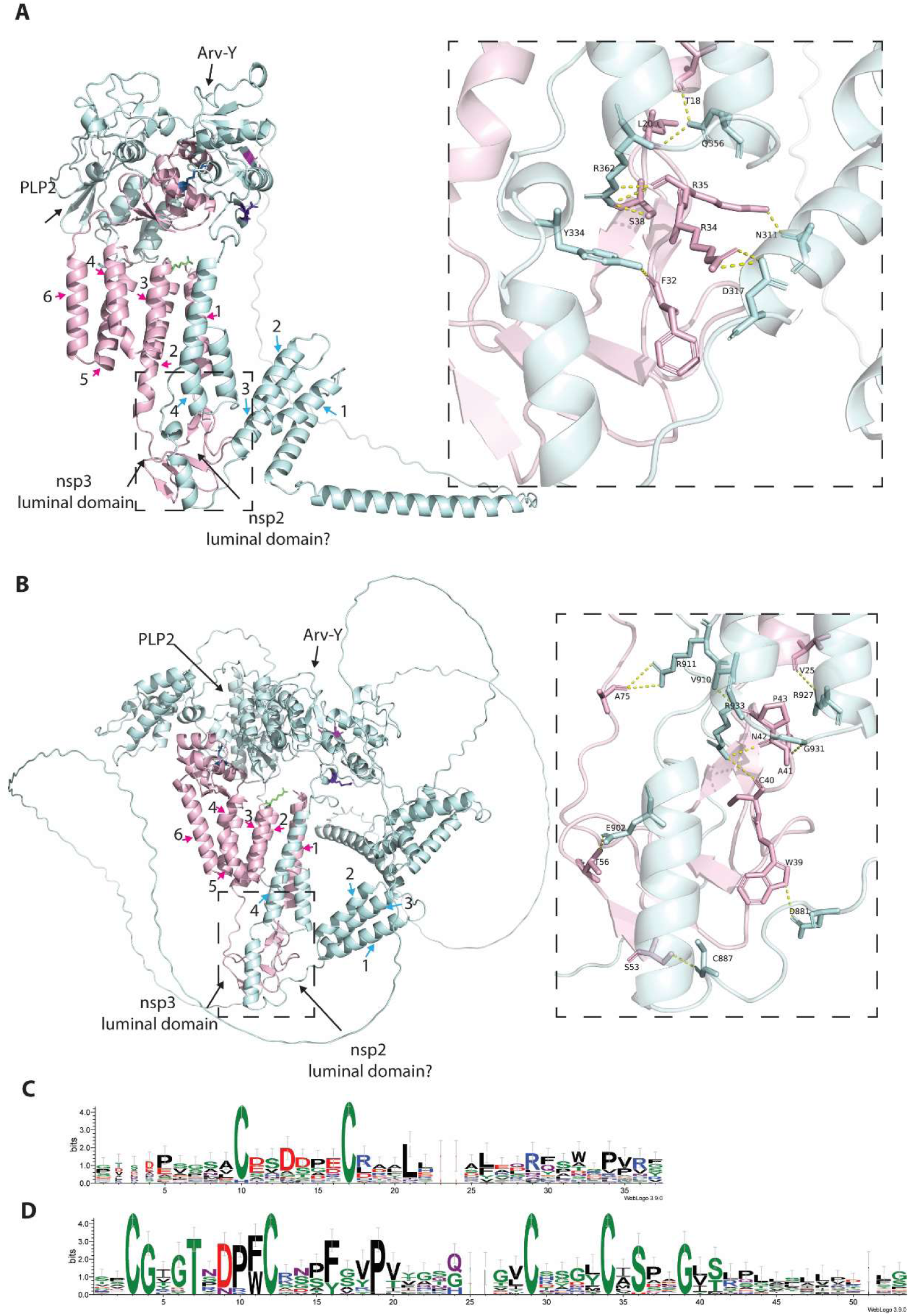
AlphaFold-based structural predictions of arterivirus nsp2-nsp3 heterodimers. AlphaFold 3 predictions of nsp2-nsp3 heterodimers for EAV (A) and PRRSV2 (B), with nsp2 shown in light cyan and nsp3 in light pink. Conserved residues highlighted in Fig. 2 are depicted using the same color scheme and shown as stick models. Predicted transmembrane α-helices (TMHs) are numbered and indicated by arrows in colors corresponding to their respective protein. In the enlarged panels, predicted interactions between residues within the highlighted regions (distance <3.5 Å) are shown as dashed lines. (C, D) Amino-acid sequence conservation within the predicted ER-luminal domains of nsp2 (C) and nsp3 (D) across all 23 arteriviruses in the MSL41 [ICTV], visualized using WebLogo3.0 (28).

To further investigate the potential architecture of the DMV pore complex, we explored whether AlphaFold-based modeling of nsp2 and nsp3 hexamers would yield structures resembling the hexameric rings that form the core of the SARS-CoV-2 nsp3-nsp4 pore complex (17). Owing to computational constraints, we modeled partial nsp2 hexamers (starting at pp1a residue 435 for EAV and 1223 for PRRSV2) together with full-length nsp3 hexamers for EAV and PRRSV2. These segments encompass the arterivirus domains postulated to correspond to the well-resolved, hexameric pore-flanking regions of the SARS-CoV-2 pore complex, namely the Arv-Y domain, the TMH2-6 segment of nsp3, and the nsp3-CTD.

For both arteriviruses, the Y1-Y2 domain structure was predicted with high confidence (pLDDT > 70), and the resulting hexameric model exhibits PAE and interface-predicted template modeling (piTM) scores that support the reliability of the inferred inter-chain interactions (Fig. S6). Strikingly, the hexameric rings formed by the Y1 domain of both EAV (Fig. 4A,C) and PRRSV2 (Fig. 4B,D) resemble the Y1-domain ring of SARS-CoV-2 nsp3, featuring a conserved sheet-loop-sheet motif lining the pore channel (17). Moreover, this arrangement fits well within the upper constriction observed in the EAV cryo-EM map (25) (Fig. 4E). Unfortunately, hexameric predictions involving the transmembrane regions of nsp2 and nsp3 lacked sufficient confidence and were inconsistent with prior data or structural expectations. AlphaFold frequently clustered TMHs in unrealistic conformations and did not account for the atypical double-membrane architecture of the vesicles in which these pore complexes reside. Additional modeling attempts using truncated sequences to generate hexamers or dodecamers did not yield structures with sufficiently high PAE and ipTM scores to permit reliable fitting into the cryo-EM density map.

**Figure 4:**
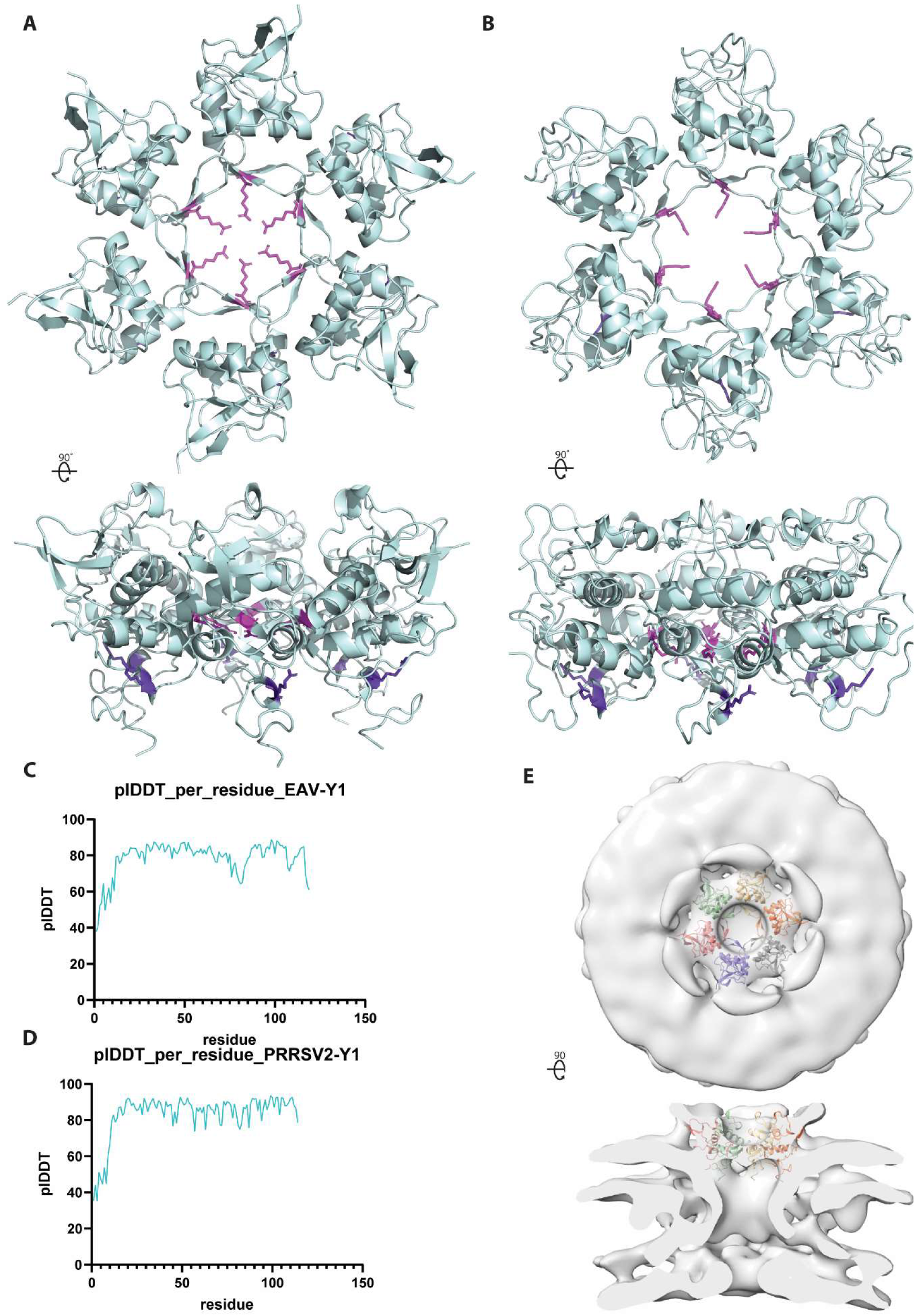
Structural predictions for the region surrounding the upper constriction of the arterivirus DMV pore channel. AlphaFold 3-based prediction of nsp2 Y1-domain hexameric rings for EAV (A) and PRRSV-2 (B), which are postulated to shape the upper constriction of the DMV pore channel. Conserved positively charged nsp2 residues are depicted as stick models in magenta (EAV R673) and purple (EAV R659). The pLDDT values were plotted for the EAV (C) and PRRSV2 (D) predictions. (E) The EAV nsp2-Y1-Y2 hexamer roughly fits within the hexagonal platform in the crown of the cryo-electron tomography-derived density map at high-threshold surface rendering of the DMV pore complex (adapted from (25), see also Fig 1F.)

In summary, monomeric modeling of EAV and PRRSV-2 nsp2-nsp3 revealed an overall topology of these transmembrane proteins that appear to mirror that of the SARS-CoV-2 nsp3-nsp4 complex. Additionally, a putative luminal domain was identified within arterivirus nsp2, which likely interacts with the luminal domain of nsp3, analogous to the nsp3-nsp4 interaction resolved in the SARS-CoV-2 pore complex. Modeling of EAV and PRRSV2 nsp2-nsp3 hexamers yielded a hexameric ring of Y1 and Y2 domains, positioned around the upper constriction site of the arterivirus pore complex. Both the architecture and predicted localization of this ring resemble those observed in the SARS-CoV-2 nsp3-nsp4 pore complex. Together, the similarities in domain organization, membrane topology, and the structural features surrounding the upper constriction site strongly suggest that arterivirus DMV pores are organized according to the same general architectural principles as their coronavirus counterparts.

### Arterivirus-wide conservation of positively-charged nsp2 and nsp3 residues predicted to face the DMV pore channel

As described above, both the SARS-CoV-2 nsp3-nsp4 pore complex and the EAV DMV pore average feature three constrictions along the pore channel (Fig. 1C,E) (17, 25). Along the coronavirus pore channel, rings of six conserved positively charged amino acid residues were identified, which face the pore channel and were implicated in RNA transport across the pore (17). To evaluate whether this feature is conserved in arteriviruses, we combined the information obtained from the nsp2 and nsp3 sequence alignments and AlphaFold modeling.

In our model, the nsp2 Arv-Y domain is the most likely candidate to line the upper constriction site (Fig. 2A-C, pink), as its position relative to the transmembrane regions parallels that of the nsp3 Y1 domain, which encompasses this constriction in the SARS-CoV-2 pore complex (Fig. 2A, pink asterisk). Additionally, the Y1-Y2 hexamer predicted by AlphaFold fitted around the upper constriction site in the EAV pore average (Fig. 4E). The middle constriction site of the SARS-CoV-2 nsp3-nsp4 pore channel (green asterisk in Fig. 2B) is surrounded by residues located between TMH2 and TMH3 of nsp4. Although precise prediction of the membrane topology of arterivirus nsp3 remains challenging, cytosolic loops connecting its TMHs represent strong candidates to occupy the corresponding second constriction of the arterivirus pore channel (Fig. 2B-C, green). The third constriction, located at the base of the channel, is lined by the nsp4-CTD in coronaviruses; accordingly, the nsp3-CTD emerges as its most plausible arterivirus equivalent (Fig. 2B-C, blue).

We next searched for conserved positively-charged residues in the nsp2 and nsp3 regions that may be involved in protein-RNA interactions relevant to RNA export through the pore. Within the nsp2 Arv-Y domain, which would face the upper constriction, two conserved positive charges were identified (EAV pp1a residues 659 and 673; Fig. 2D, purple and light-pink arrow). Regarding the predicted middle constriction, 13 out of 23 arterivirus nsp3 sequences contain a cytosolic loop harboring a positively-charged residue (EAV pp1a residue 920; Fig. 2D, green arrow). In most of the remaining sequences, this position is occupied by an alternative residue with properties associated with protein-RNA interactions, such as aromatic residues (Fig. 2D, Fig. S4). Finally, within the nsp3-CTD, which likely forms the lower constriction, a single positively-charged residue is conserved among arteriviruses (EAV pp1a residue 1043; blue arrow).

Of the two conserved positively-charged EAV nsp2 residues in our model, the second (R673; Fig. 2D; light pink) is predicted to face the pore channel, whereas the first (R659; Fig. 2D; purple) would be oriented towards the membrane interface (Fig. 4A-B). While the hexameric predictions involving the transmembrane regions of nsp2 and nsp3 lacked sufficient confidence, AlphaFold predictions for nsp3 supported the positioning of the identified conserved positively-charged residues along the channel (Fig. 2). The residue between TMH2 and TMH3 (EAV R920; Fig. 2D,3A; green) maps within the cytosolic loop connecting these helices, while the second conserved residue (EAV R1043; Fig. 2D, 3A; blue) is located on an exposed surface of the nsp3-CTD.

In conclusion, sequence comparisons and AlphaFold-based modeling of EAV and PRRSV2 nsp2 and nsp3 support the notion that the overall organization of the arterivirus DMV pore complex is broadly similar to that of the SARS-CoV-2 nsp3-nsp4 pore. Of the four conserved positively-charged residues initially identified, three (R673, R920, and R1043 in EAV) were consistently predicted to face the pore channel, where they could potentially contribute to RNA translocation. In contrast, R659 would be oriented towards the outer DMV membrane. Next, all four conserved arginine residues were subjected to targeted mutagenesis to experimentally validate these structural insights and assess their functional significance.

### Conserved positively charged residues at predicted EAV pore channel constrictions are essential for replication

The four conserved positively-charged residues (R659 and R673 in nsp2, R920 and R1043 in nsp3; Fig. 2D) were targeted by reverse genetics using EAV full-length cDNA clone pEAN551 (32). Each arginine was replaced either with lysine, a conservative positively-charged substitution, or with alanine, a small neutral amino acid. Virus mutants were launched by transfecting in vitro-transcribed full-length RNA into BHK-21 cells, which were subsequently monitored for the development of cytopathic effects. The parental recombinant wild-type virus (rWT) was included as a positive control. At 10, 24, 48, and 120 h post-transfection, transfected cells were fixed for immunofluorescence microscopy and stained with antibodies recognizing a nonstructural protein (nsp3) and a structural protein (GP_5_). Furthermore, culture supernatants were harvested for plaque assays and sequence analysis of progeny virus. Virus mutants were deemed non-viable if two independent attempts to recover them failed to yield cells positive for viral proteins and/or detectable infectious progeny.

Transfections with all four mutants carrying conservative Arg-to-Lys substitutions, as well as the wt control virus, induced cytopathic effects and yielded detectable levels of infectious progeny (Fig. 5A). In contrast, none of the Arg-to-Ala substitution mutants produced a detectable amount of infectious progeny (Fig. 5A) or showed positive staining for viral proteins by immunofluorescence microscopy (data not shown), indicating that replacement of the arginine residues with a neutral amino acid is lethal to virus replication. Across all harvesting time points, transfection with mutant R920K yielded wt virus-like plaques and progeny titers (Fig. 5A-B). In contrast, at 24 and 48 h post-transfection, the R659K and R1043K samples showed severely reduced progeny titers, although titers approached those of the wt control by 120 h post-transfection. For both these mutants, we noticed a mixed plaque phenotype with an increased proportion of small(er) plaques compared to the wild-type control. Finally, transfection of mutant R673K resulted in a measurable progeny titer only by 120 h post-launch (Fig. 5A-B), with a heterogeneous plaque phenotype characterized by generally smaller plaques.

**Figure 5:**
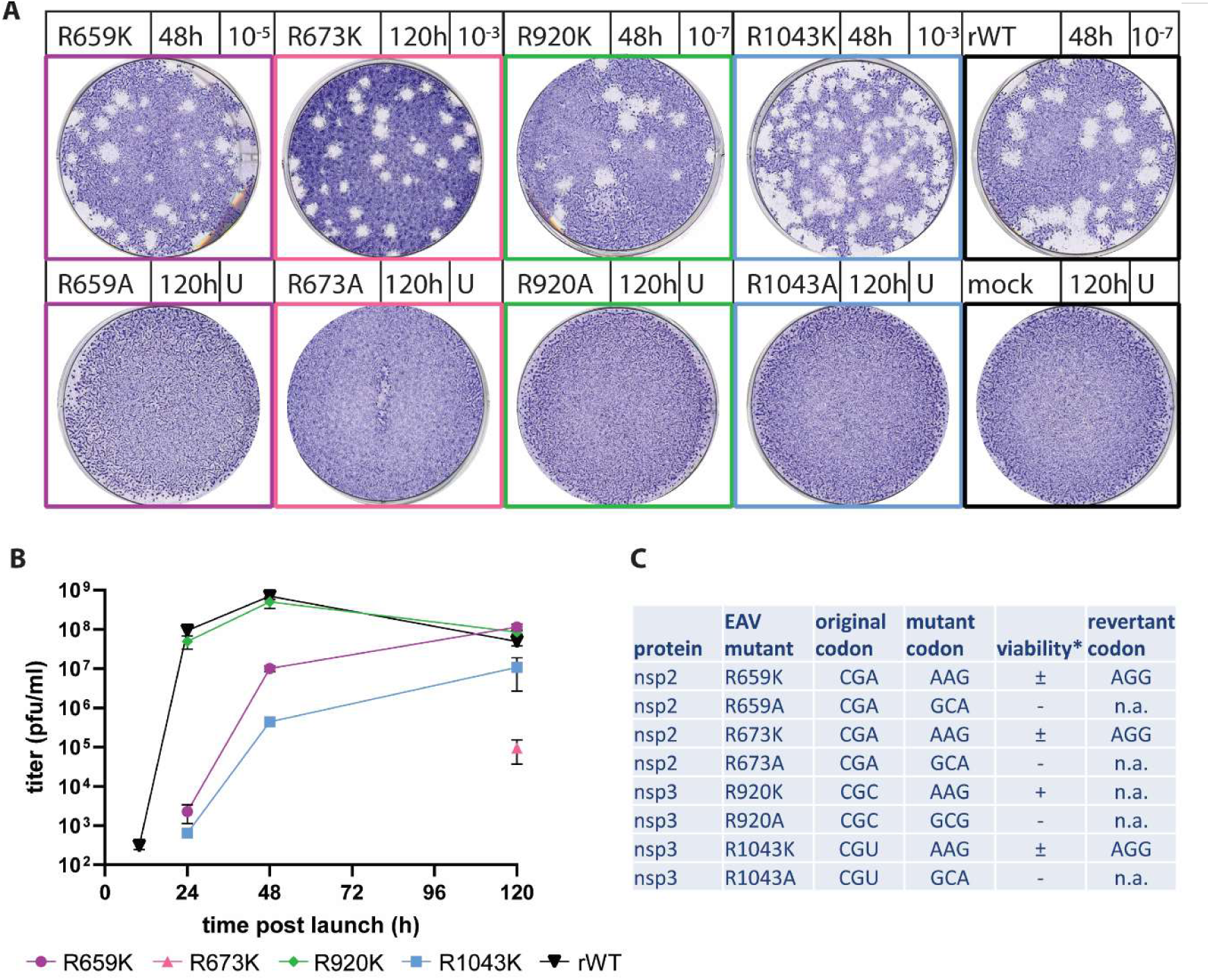
Replacement of conserved arginine residues in EAV nsp2 and nsp3 with alanine is lethal. (A) Plaque assays of transfection samples of EAV mutants launched in BHK-21 cells; the mutation, time of harvest, and sample dilution used (0.5 ml inoculum per well) are indicated above each image; U, undiluted sample. (B) For transfections of viable mutants, the amount of infectious progeny released into the supernatant was monitored by plaque assay up to 120 h post-launch; data are represented as mean ± SD with n=3. (C) Summary of the viability and replication of the nsp2 and nsp3 mutants; –, non-viable; ± reversion detected; +, similar to wt virus.

The aberrant growth characteristics and plaque phenotype of EAV-R673K, EAV-R659K, and EAV-R1043K suggested rapid post-launch evolution. This was confirmed by RT-PCR amplification and sequence analysis of the nsp2-nsp3-coding region, which revealed that reversion occurred between 2 and 5 days post-transfection. In each case, the restored arginine codon (AGG) differed from the original codon in the EAV genome (CGN), thereby excluding the possibility of contamination with wild-type virus (Fig. 5C). In contrast, evidence of reversion was not observed for EAV-R920K, highlighting a unique tolerance for lysine at this position.

In summary, our results strongly suggest that a positive charge is essential for EAV replication at each of the four conserved positions in nsp2 and nsp3, as substitutions that eliminated these charges were lethal. Although lysine could apparently functionally substitute for arginine at these positions to varying degrees, arginine appears to be the preferred residue at three of the four sites. The replication of the corresponding Arg-to-Lys mutants appeared to be seriously impaired, and the viral progeny titers measured at 2 to 5 days post-transfection should largely be attributed to reversion.

### Substitution of conserved positively-charged residues in EAV nsp2 and nsp3 does not affect processing of the nsp2/3 junction or the nsp2-nsp3 interaction

To further investigate the impact of substituting the conserved arginine residues in EAV nsp2 and nsp3, the same eight mutations, including the four alanine replacements that were lethal in the context of virus replication, were introduced into the previously described expression vector pcDNA-HAnsp23GFP (20). This construct directs the expression of a self-cleaving polyprotein consisting of an N-terminally HA-tagged EAV nsp2 and a C-terminally GFP-tagged nsp3. Expression of these two proteins is sufficient to induce the formation of paired membranes and DMVs (20), making this system well-suited to study the role of the conserved arginines outside the context of EAV-infected cells.

We first analyzed PLP2-mediated processing of the nsp2/3 junction and the interaction between nsp2 and nsp3, both of which are essential for viral replication (21, 32). Prior to cell lysis at 24 h post-transfection, the GFP signal associated with the C-terminally tagged nsp3 was used to confirm comparable transfection efficiencies of H1299 cells for all constructs. Subsequently, protein lysates were prepared, and nsp3-GFP, together with any interacting HA-nsp2, was pulled down using GFP-catcher beads containing a covalently bound anti-GFP nanobody. Total HA-nsp2 levels in the transfected cell lysates were monitored using an antibody recognizing the C-terminal region of nsp2.

Expression of the wt HA-nsp2-nsp3-GFP resulted in complete nsp2/3 cleavage and efficient nsp2-nsp3 interaction. As anticipated, replacement of one of PLP2’s catalytic residues (H332Y; (49)) abolished cleavage, yielding only the full-length HA-nsp2-3-GFP (113 kDa; Fig. 6). None of the eight arginine substitutions substantially affected processing of the nsp2/3 site, although a faint band corresponding to unprocessed HA-nsp2-3GFP was detected for the R659A and R920A mutants (Fig. 6). Consistent with previous observations (21), pull-down of nsp2 together with nsp3 was readily detectable for the wt control. Importantly, for all mutants, HA-nsp2 pull-down was successful and, when accounting for HA-nsp2 levels in the input material, pull-down efficiency appeared comparable to that observed for the wt proteins (Fig. 6). In conclusion, the arginine substitutions ins nsp2 and nsp3 did not interfere with efficient nsp2/3 cleavage or with the formation of the stable nsp2-nsp3 interaction that was previously documented in EAV-infected cell lysates (21).

**Figure 6:**
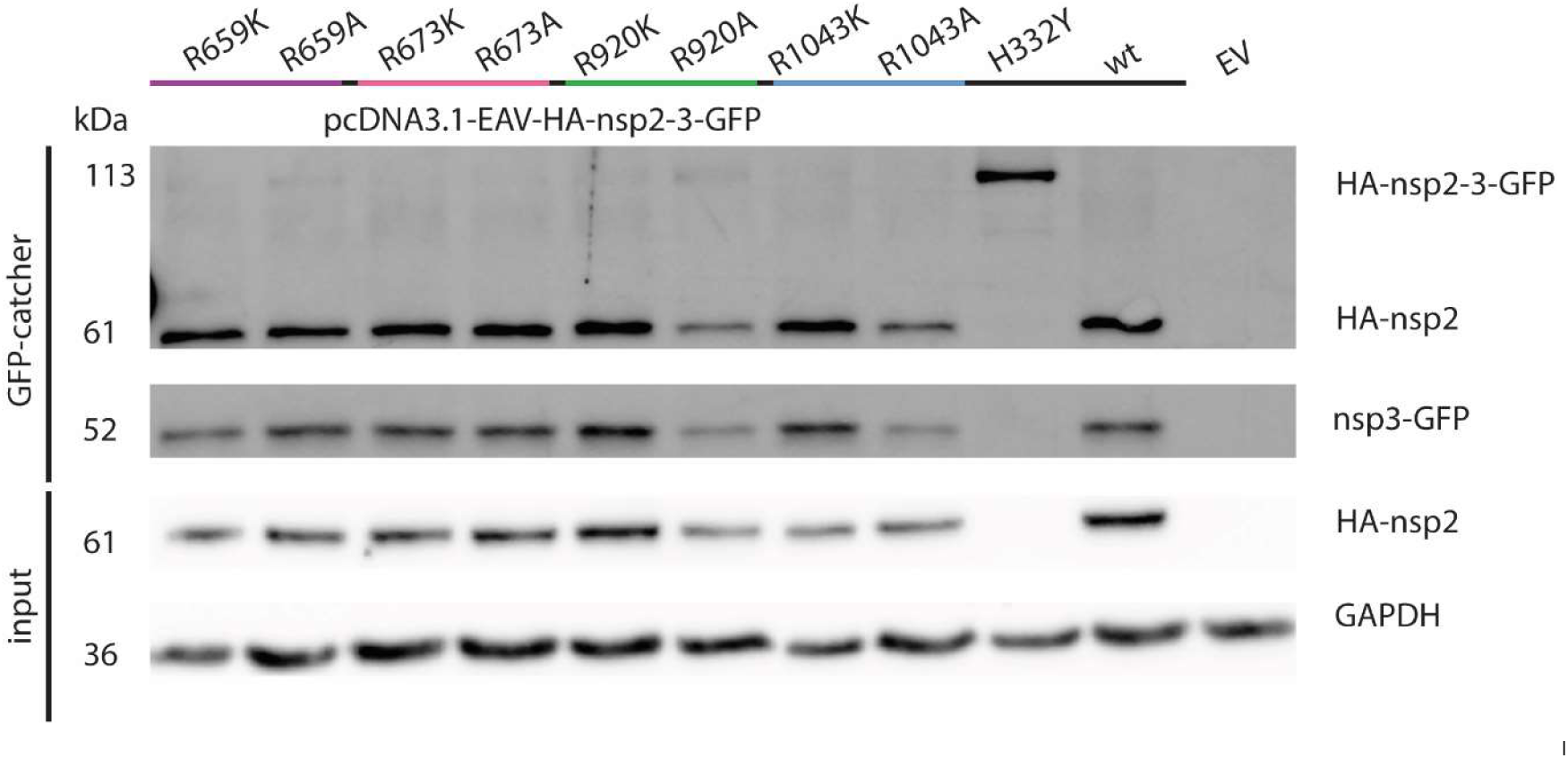
Substitution of conserved arginine residues in EAV nsp2 and nsp3 does not affect nsp2/3 cleavage or nsp2-nsp3 interaction. Analysis of lysates from cells expressing self-cleaving (wt or mutant) HA-nsp2-nsp3-GFP polyprotein, to assess EAV nsp2/nsp3 cleavage and the interaction of EAV nsp2 and nsp3. GFP-catcher beads were used to pull down nsp3-GFP and any associated HA-nsp2. Input samples and GFP-catcher bead elution samples were separated by 10% SDS-PAGE. Protein bands were visualized by Western blotting, using a rabbit antiserum recognizing GFP and the CTD of EAV nsp2, to assess polyprotein processing and the nsp2-nsp3 interaction, as reflected by HA-nsp2 pull down with nsp3-GFP. Wild-type and PLP2 catalytic mutant (H332Y) constructs were included as controls.

### DMV formation is maintained after substitution of pore channel-facing arginine residues but does depend on R659 in nsp2

As nsp2 and nsp3 constitute the minimal machinery required for DMV and DMV pore formation (10, 16, 24, 25), we examined whether substitution of the conserved arginine residues in these proteins affected this process. To this end, transmission electron microscopy (TEM) was used to analyze cells transfected with the mutant expression constructs described above. In cells expressing wt HA-nsp2-3-GFP, DMVs were the predominant membrane structures observed, although putative intermediates in their biogenesis, such as zippered ER, were also detected, consistent with previously published work (Fig. 7) (20, 25).

**Figure 7:**
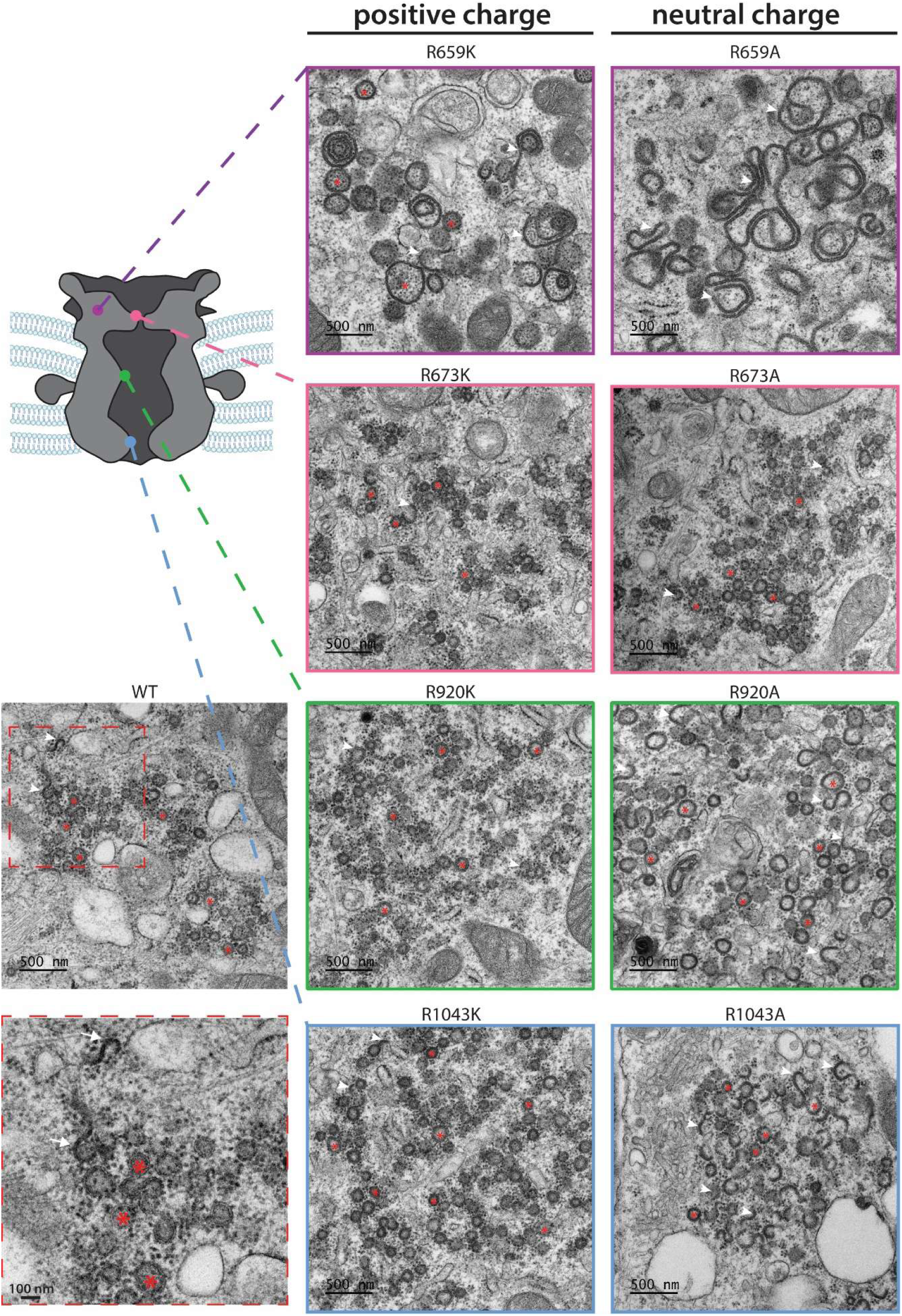
EM analysis of membrane rearrangements induced by expression of EAV nsp2-nsp3 mutants carrying substitutions of conserved arginine residues. TEM images revealed DMV formation (examples marked by red asterisks) for all nsp2-nsp3 mutants except R659A. In addition to fully formed DMVs, various putative intermediates of DMV biogenesis were visible (examples highlighted by white arrowheads). A magnified view of representative DMVs and membranous intermediates present in cells expressing the wt protein is marked with a red-dashed box. The predicted locations of the mutated residues are indicated in the accompanying schematic representation of the EAV pore complex, using the previously assigned color scheme.

For mutants in which the three arginine residues predicted to face the pore channel were replaced (Fig. 2C), DMV formation was readily observed for both positive-to-positive and positive-to-neutral charge substitutions (Fig. 7, red asterisks). In all specimens, paired membrane structures were also detected (Fig 7, white arrowheads), consistent with ongoing or incomplete DMV biogenesis. Notably, such paired membranes (white arrowheads) were more frequently observed in cells expressing the R920A and R1043A mutants, whereas the structures in the cells expressing their positive-to-positive charge mutation counterparts more closely resembled those generated by the wt control. Taken together, all mutants retained the ability to induce DMVs that were morphologically similar to those induced by wt nsp2-nsp3, which suggests that the conserved arginine residues proposed to face the pore channel play a limited, if any, role in DMV formation itself. This finding is consistent with their proposed function in processes occurring after DMV and pore biogenesis, such as viral RNA export through the pore complex.

In contrast, DMV formation appeared strongly impaired upon substitution of residue R659, which is predicted to face the outer DMV membrane (Fig 4A,B) and therefore is unlikely to be directly involved in RNA export (Fig. 7, purple). Introduction of a positive-to-neutral substitution at this position (R659A) completely abolished detectable DMV formation; instead, extensive regions of zippered ER were present. This double-membrane intermediate of DMV biogenesis is thought to arise from nsp2-nsp3 interactions, which do remain intact in the R659A mutant according to the pull-down results presented above (Fig. 6). Replacement of R659 with lysine (R659K) did not fully restore DMV formation, although occasional DMV-like structures —substantially larger than those induced by wt nsp2-nsp3— could be detected. Collectively, these observations indicate that a positive charge at position 659 in nsp2 is essential for DMV formation and that arginine is preferred over lysine at this site. Importantly, this requirement appears to be independent of the nsp2-nsp3 interaction that mediates membrane pairing. Together, our data suggest that the conserved arginine residues in nsp2 and nsp3 play separable roles, with pore-facing residues likely acting downstream of DMV formation, whereas R659 is critically involved in the membrane remodeling events required for DMV biogenesis itself.

## Discussion

Nidoviruses share several defining features in genome organization and replication strategy and may also employ common mechanisms for the formation of replication organelles in the cytoplasm of infected cells. However, studies of this process have so far been largely limited to corona- and arteriviruses. One of the most significant recent advances is the identification of a conserved molecular pore complex spanning the double membrane of the DMVs that constitute the central scaffold of the replication organelles induced by arteri- and coronaviruses (15–17, 25). In both families, DMV pore complexes are formed by two viral nonstructural proteins that contain transmembrane domains. These proteins also act as key drivers of membrane pairing and DMV biogenesis, independently of their role in pore formation (16, 17, 21, 24, 25). Notably, while interactions between the ER-luminal domains of these proteins are thought to be required for membrane pairing, additional, as-yet-uncharacterized interactions – either between the two proteins or among copies of the same protein – are likely essential for efficient DMV and/or pore formation.

Due to the low-throughput nature of the FIB-milling and cryo-ET workflow, we have thus far been unable to obtain a structure of the arterivirus DMV pore at sufficient resolution to perform de novo model building (25). Nevertheless, within the constraints imposed by the currently available low-resolution (20 Å) structure, the EAV pore appears to exhibit an architecture strikingly similar to that of its coronavirus counterpart. Both are centered around a putative RNA export channel whose diameter varies along its length. Despite its substantially smaller dimensions, the arterivirus complex is likewise thought to consist of two times six copies each of nsp2 and nsp3, arranged into four stacked hexameric rings (25) (Fig. 1). Like coronavirus nsp3 and nsp4, arterivirus nsp2 and nsp3 are proposed to interact via their luminal domains within the DMV intermembrane space, forming a belt-like structure with pseudo-12-fold symmetry that encircles the complex (25). The estimated mass (∼1.4 MDa) of the EAV pore complex is consistent with this interpretation and underpins the structural analysis presented in Figures 2 to 4 of this study.

Alignment of nsp2 and nsp3 amino acid sequences across all established arteriviruses species highlighted multiple well-conserved domains. Many of these, including the PLP2 protease, the predicted TMHs, and conserved cysteine-containing motifs, have been described previously (20, 32). While this manuscript was in preparation, the structural conservation of the Y1, Y2, and Y4-like motifs within the Arv-Y domain of arterivirus nsp2 was reported (37). Notably, related motifs are present in SARS-CoV-2 nsp3 and nsp4, providing a framework for developing a preliminary model of the architecture and domain organization of the arterivirus pore complex (Fig. 2D). Identifying conserved structural and functional features shared between distantly related nidovirus families may facilitate the discovery of novel broadly acting antivirals targeting RO formation or RNA export.

Using sequence comparison in combination with the current model of the EAV pore complex (25), supplemented with AlphaFold predictions, we identified four conserved positively-charged residues of interest. Three of these were predicted to face the pore channel, while R659 in nsp2 appears to map to the membrane-facing surface of the pore (Fig. 4A-B). This residue lies adjacent to the recently described zinc-binding domain in nsp2 (37), suggesting a function distinct from that of the pore-lining arginine residues. To assess the functional relevance of these residues, each was mutated either to retain a positive charge (Arg-to-Lys) or to eliminate it (Arg-to-Ala), and the viability of the resulting virus mutants was evaluated (Fig. 5).

In line with the predicted importance of maintaining a positive charge at these sites, all Arg-to-Lys mutants could be launched, whereas none of the Arg-to-Ala mutants showed evidence of replication, possibly due to the disruption of essential interactions between the DMV pore channel and viral RNA. Notably, only EAV-R920K displayed wt-like replication, while the launching of the remaining viable mutants produced initially smaller plaques and substantially reduced progeny titers (Fig. 5 A-B). For these crippled Arg-to-Lys mutants, sequence analysis of progeny harvested at later time points confirmed rapid reversion to arginine, likely facilitated by the requirement for only a single nucleotide change (AAG→AGG). This finding indicates a strong preference for arginine at two of the three putative constriction sites, which is unlikely due to steric constraints, given that arginine is larger and more rigid than lysine. Instead, arginine offers a greater capacity for polar interactions and salt-bridge formation (50). Consistent with this notion, arginine residues play central roles in a wide range of RNA-protein interactions, including those involving other viral proteins (18, 51–53). In some systems, arginine residues are specifically required for RNA translocation through dynamic RNA-protein interaction cascades (52).

Beyond the well-described general role of arginine residues in protein-RNA interactions, several domains in SARS-CoV-2 proteins, including the conserved nsp3 Y1 and CoV-Y domains, have been reported to exhibit specific affinity for the 5’ end of the viral genome (54). Similar interactions may facilitate the transit of viral RNA through the DMV pore channel. A gradient of increasing affinity between successive constriction sites and the RNA could provide a passive transport mechanism, particularly given that the viral RNA polymerase has not yet been detected as an integral component of the pore complex and thus may not actively drive RNA extrusion. On the other hand, small asymmetric masses have been detected at the base of the pore complex in cells infected with the coronavirus mouse hepatitis virus and EAV (15, 25). These require further investigation, but could represent part of the viral RNA polymerase complex and may contribute to RNA export. Additionally, in an expression system, the heterodimerization of nsp2-nsp3 in PRSSV2 was recently shown to regulate binding of the RdRp-containing nsp9 subunit. Subsequently, mutagenesis of residues crucial for this interaction proved to be lethal for the virus (48). Nevertheless, it remains to be experimentally demonstrated that viral RNA indeed transits through the DMV pore and, if so, by what precise mechanism this process is achieved.

To further investigate the role of conserved arginine residues in nsp2 and nsp3 in arterivirus DMV formation, mutations were introduced into an nsp2-3 expression system, thus bypassing the need for viral replication. In this context, neither nsp2/nsp3 cleavage nor the interaction between nsp2 and nsp3 - processes analogous to those that are crucial for coronavirus DMV and pore formation (16, 17) - were impaired in any of the mutants (Fig. 6). TEM analysis of cells expressing these constructs showed that substitution of residues predicted to line the pore-channel did not hinder DMV formation. In contrast, mutations at R659, which is not predicted to face the pore channel, had a profound impact on DMV biogenesis. The R659A mutant failed to form DMVs and instead accumulated membrane rearrangement intermediates, such as zippered ER. The conservative R659K mutation partially restored DMV formation but resulted in enlarged DMVs, along with an increased abundance of putative intermediates of DMV biogenesis (Fig. 7).

The abundant zippered ER observed upon substitution of R659 (Fig. 7) indicates that the resulting defect in DMV formation is independent of membrane pairing. This is consistent with the preservation of the nsp2-nsp3 interactions (Fig. 6), which are thought to be essential for membrane pairing. This suggests a role for this residue in protein folding, oligomerization, or additional protein-protein interactions required for DMV maturation. Strikingly, mutation of this single residue, within a domain lacking any known enzymatic function, was sufficient to completely abrogate DMV formation, providing a plausible explanation for the lethal impact of this substitution when introduced into the viral replicase (Fig. 5). Notably, R659 is positioned adjacent to a conserved zinc-binding motif, which has been suggested to locally destabilize the membrane and, upon nsp2 oligomerization, release zinc ions to liberate the Y1 domain for pore formation (37). Such conformational and membrane-associated changes could facilitate critical interactions during the transition from early membrane remodeling events to DMV maturation and pore formation, a process in which the positive charge of the nearby R659 residue may play an important structural or regulatory role. Ultimately, high-resolution structural information on the arterivirus DMV pore complex will be required to elucidate the precise interactions governing DMV biogenesis, pore assembly, and RNA export.

## Acknowledgments

We thank Stanley Fronik for sharing the EAV pore averages prior to publication and Jason Perry for helpful discussion regarding the AlphaFold predictions. In addition, we would like to thank many LUMC colleagues for helpful discussions. This study was supported in part through the European Union’s Horizon Europe research and innovation program under grant agreement 101137229 (PANVIPREP) and the Netherlands Organization for Scientific Research under grant agreement NWO-OCENW.M21339.

## Data availability

All data generated or analyzed are included in this article and its supplementary material.

